# Dietary protein source mediates colitis pathogenesis through bacterial modulation of bile acids

**DOI:** 10.1101/2025.01.24.634824

**Authors:** Simon M. Gray, Michael C. Wood, Samantha C. Mulkeen, Sunjida Ahmed, Shrey D. Thaker, Bo Chen, William R. Sander, Vladimira Bibeva, Xiaoyue Zhang, Jie Yang, Jeremy W. Herzog, Shiying Zhang, Belgin Dogan, Kenneth W. Simpson, R. Balfour Sartor, David C. Montrose

**Author notes:** Authors contributed equally (Simon M. Gray, Michael C. Wood). **Corresponding Author**: David C. Montrose, Department of Pathology, Renaissance School of Medicine, Stony Brook University, MART Building, 9M-0816, Lauterbur Dr., Stony Brook, NY 11794. **Declaration of interests:** D.C.M. has consulting arrangements with Gleipnir Medical. The authors have no other competing interests relevant to this study.

## Abstract

Evidence-based dietary recommendations for individuals with inflammatory bowel diseases (IBD) are limited. Red meat consumption is associated with increased IBD incidence and relapse in patients, suggesting that switching to a plant-based diet may limit gut inflammation. However, the mechanisms underlying the differential effects of these diets remain poorly understood. Feeding diets containing plant- or animal-derived proteins to murine colitis models revealed that mice given a beef protein (BP) diet exhibited the most severe colitis, while mice fed pea protein (PP) developed mild inflammation. The colitis-promoting effects of BP were microbially-mediated as determined by bacterial elimination or depletion and microbiota transplant studies. In the absence of colitis, BP-feeding reduced abundance of *Lactobacillus johnsonii* and *Turicibacter sanguinis* and expanded *Akkermansia muciniphila*, which localized to the mucus in association with decreased mucus thickness and quality. BP-fed mice had elevated primary and conjugated fecal bile acids (BAs), and taurocholic acid administration to PP-fed mice worsened colitis. Dietary psyllium protected against BP-mediated inflammation, restored BA-modulating commensals and normalized BA ratios. Collectively, these data suggest that the protein component of red meat may be responsible, in part, for the colitis-promoting effects of this food source and provide insight into dietary factors that may influence IBD severity.

## Main Text

Diet influences the pathogenesis and natural history of inflammatory bowel diseases (IBD), disorders of chronic intestinal inflammation caused by dysregulated host immune responses to resident intestinal microbes in genetically susceptible hosts^1–3^. Despite these findings, limited evidence-based dietary recommendations exist to reduce the incidence and severity of IBD^4–6^. For multiple decades IBD incidence has increased in the U.S. and globally, including in regions with previously low rates^7–9^. Increased animal-based protein consumption, including red meat, correlates with rising IBD incidence, suggesting a link between animal protein consumption and IBD^10, 11^. However, the mechanisms by which specific dietary protein sources promote gut inflammation remains unclear.

Diet regulates gut homeostasis and inflammation, in part, by shaping the composition, function, and metabolic activity of the gut microbiota^2, 12–21^. For example, reducing dietary fiber or increasing fructose consumption in mice enriches mucus-digesting resident bacteria, such as *Akkermansia muciniphila*^12, 17–19, 21^. Further, consuming high fructose or dietary milk fat limits the deconjugation of bile acids (BAs) (small molecules synthesized by the liver and modified by intestinal microbes), resulting in increased levels of conjugated BAs (CBAs) and exacerbated experimental colitis^17, 22^. Notably, patients with active IBD have higher levels of primary and CBAs and depleted secondary unconjugated BAs, relative to healthy controls^23–25^. Moreover, primary CBAs, which are metabolized by microbial enzymes to unconjugated secondary BAs, exacerbate inflammation in experimental colitis models^22, 24, 26^. However, the role of dietary protein source in these complex interactions and development of colonic inflammation is poorly understood.

To investigate the impact of dietary protein source on experimental colitis severity, five isocaloric synthetic diets containing protein isolates from beef, egg whites, casein, soy or pea (**Table S1**) were fed to wild-type (WT) specific pathogen-free (SPF) C57BL/6J mice during dextran sodium sulfate (DSS)-mediated induction of colitis. Mice fed beef protein (BP) diet developed the most severe colitis while those fed pea protein (PP) diet developed the least severe colitis; egg whites, casein, or soy protein diets induced intermediate colitis severity (**Fig. 1A-D**). BP feeding resulted in more severe colitis in a T-cell mediated colitis model using ex germ-free (GF) *Il10*^−/−^ mice colonized with mouse-adapted pooled human IBD microbiota (IMM-HM2), to induce human IBD microbiota-associated colitis^27^, compared to feeding a PP diet (**Fig. 1E-G)**. This was characterized by greater weight loss, more severe histologic inflammation and higher expression of colonic pro-inflammatory cytokines (**Fig. 1E-G)**. Similar findings were made in a second model of human IBD microbiota-associated colitis (IMM-g2)^27^ (**Fig. 1H-J**). To evaluate whether BP promotes colonic inflammation in the absence of a colitis inducer (i.e. DSS or IBD patient-derived stool), WT SPF mice were fed standard chow, PP diet, or BP diet for 8 weeks and monitored, which showed no significant differences in DAI or colonic pro-inflammatory cytokine expression **(Fig. S1A-B**). Collectively, these data demonstrate that the protein component of beef worsens colonic inflammation in multiple experimental colitis models.

**Figure 1.**
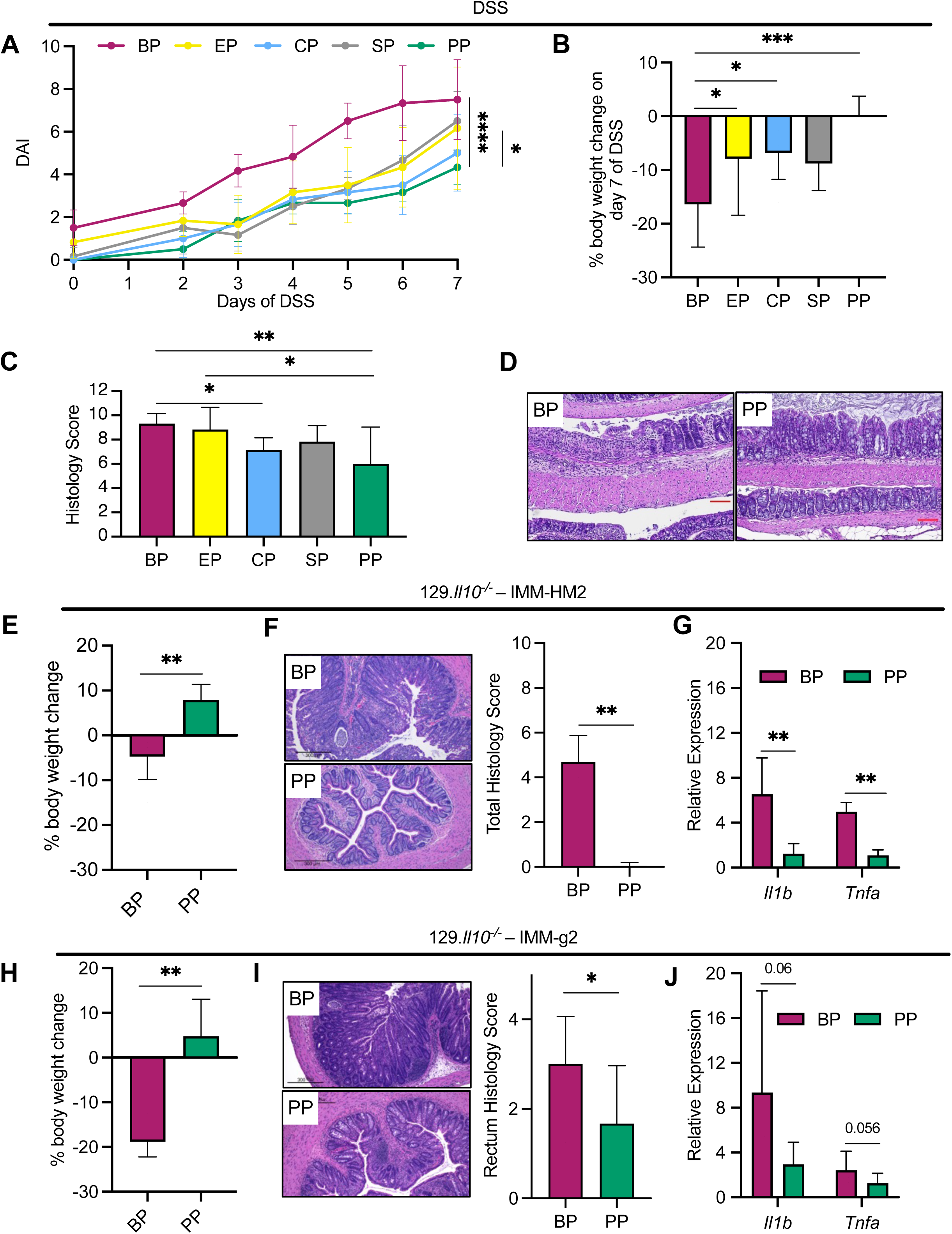
Dietary beef protein worsens murine colitis compared to other protein sources. A-D, C57BL/6 mice were fed isocaloric diets containing protein isolate derived from beef (BP), soy (SP), egg whites (EP), casein (CP), or pea (PP) for 7 days, followed by administration of 1% DSS for 7 days, while being continued on the respective diets. Disease activity index (DAI) was measured during DSS exposure (A), percent of body weight change relative to day 0 of DSS exposure was calculated following 7 days of DSS administration (B) and histologic score was assessed in colons on day 7 of DSS administration (C). n = 6 mice per group. Representative histologic images of colons from BP or PP-fed mice are shown (D). E-G, germ-free *Il10^−/−^* mice were inoculated with mouse-adapted (MA) pooled human IBD patient fecal microbiota (IMM-HM2) then fed diets containing BP or PP for 14 days. Weight change relative to day 0 was measured (E), representative histologic images of colons are shown (left side) and total histology score was assessed (right side) (F), and relative gene expression of cytokines were measured in colons (G), on day 14. n = 7-8 mice per group. H-J, germ-free *Il10^−/−^* mice were inoculated with a second MA pooled human IBD patient fecal microbiota (IMM-g2) then fed BP or PP diets for 8 days. Weight changes relative to day 0 were measured (H), representative histologic images of colons are shown (left side) and rectal histology score was assessed (right side) (I), and relative gene expression of cytokines were measured in colons (J), on day 8. n = 4-6 mice per group. Data representative of 3 (E-G), and 3 (H-J) independent experiments. *P<0.05; **P<0.01; ***P<0.001; ****P<0.0001. Bar plots show mean and standard deviation.

We next tested whether protein source modulates colitis severity, in part, through gut resident bacteria. First, WT SPF mice were fed PP diet, BP diet, or BP diet in combination with broad-spectrum antibiotics in drinking water to globally reduce gut bacteria, then challenged with DSS while continuing the respective diets and antibiotics. Antibiotic treatment reduced BP diet-mediated DSS colitis severity to, or below, PP diet-mediated levels (**Fig. 2A-C)**. An ~500-fold depletion in total bacterial abundance in feces of mice given antibiotics was confirmed by qRT-PCR. Next, GF *Il10*^−/−^ mice were fed BP or PP diets for 14 days and assessed for colonic inflammation by multiple parameters, which revealed no overt colitis in either group (**Fig. 2D-F**). GF status in both groups was confirmed by anaerobic culture of feces (data not shown). Lastly, we performed a fecal microbial transplant phenotype transfer experiment. Here, feces from WT SPF mice fed BP or PP diets for 1 week were transplanted to GF WT mice fed standard rodent chow and challenged with DSS for 10 days. Measurements of disease activity index (DAI), colon length and histologic score showed that mice gavaged with fecal slurry from BP-fed mice developed more severe colitis than mice administered feces from PP-fed mice (**Fig. 2G-I**). Together, these data demonstrate that BP exacerbates colitis through the diet-altered microbiota.

**Figure 2.**
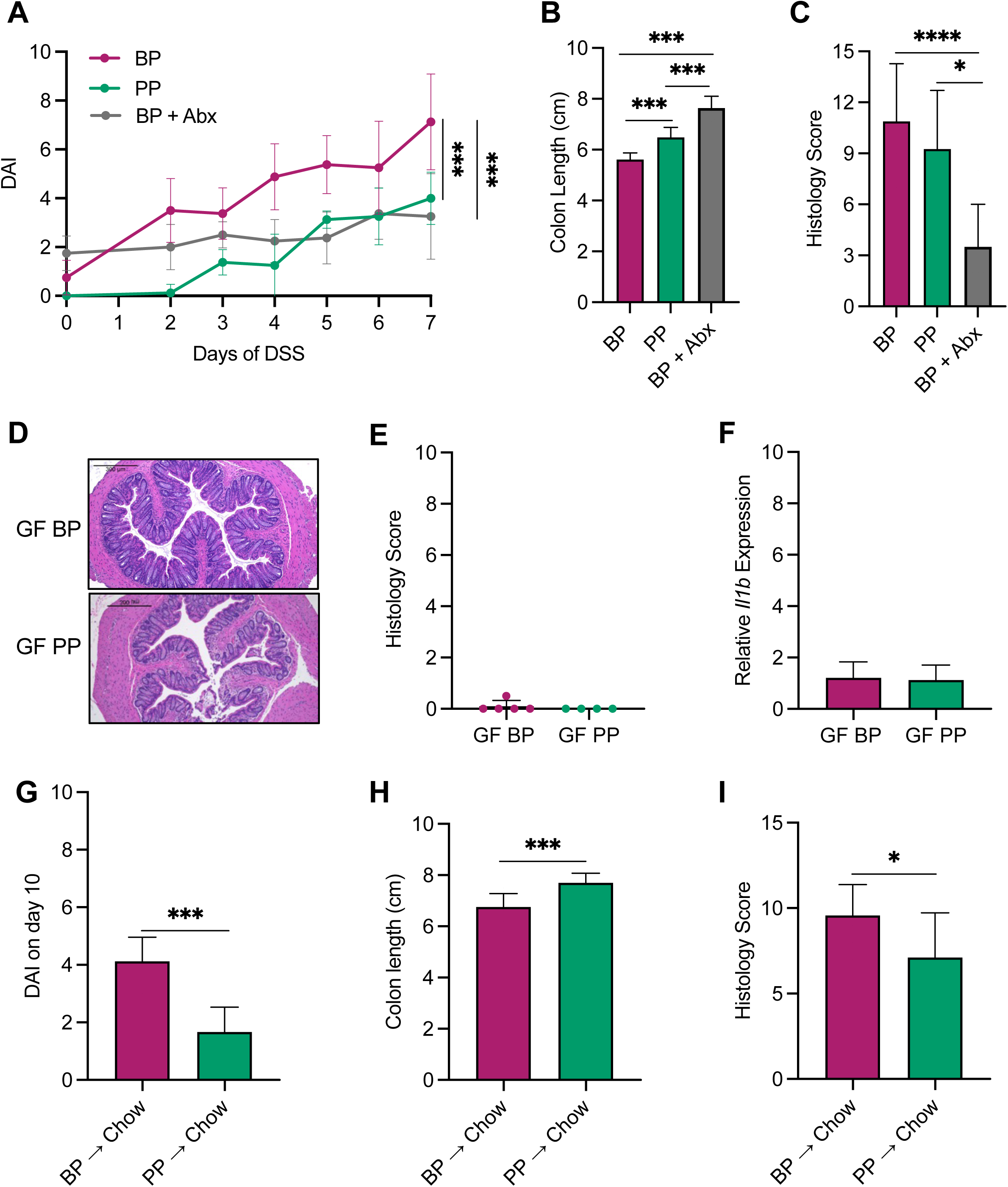
Gut bacteria are causally linked to beef protein-worsening of colitis. A-C, C57BL/6J mice were fed beef protein (BP) or pea protein (PP)-containing diets or beef protein diet and administered a five-antibiotic cocktail (ampicillin, gentamicin, metronidazole, neomycin, vancomycin^17^) in drinking water for 14 days (BP + Abx), then all mice were given 1% DSS in drinking water for 7 days while being continued on control or antibiotics-containing water. Disease Activity Index (DAI) was measured during DSS exposure (A), colon length was measured on day 7 (B) and histologic inflammation of colons was assessed on day 7 (C). n = 8 mice per group. D-F, germ-free *Il10^−/−^* mice were fed sterile BP or PP diets for 14 days. Representative histologic images of colons are shown (D), histology score was assessed (E) and relative gene expression of *Il1b* was measured in colons (F), on day 14. n = 4-5 mice per group. G-I, germ-free wild-type mice were inoculated with colonic luminal fecal contents of specific pathogen-free mice fed BP or PP diets for 7 days. Recipient mice were then fed standard chow and administered 1% DSS for 7 days and DAI was calculated (G), colon length was measured (H) and histologic inflammation of colons was assessed (I), on day 10. n = 9-10 per group; data are representative of 2 independent experiments. *P<0.05; **P<0.01; ***P<0.001; ****P<0.0001. Bar plots show mean and standard deviation.

Since the dietary protein-altered microbiota mediates colitis severity, bacterial community composition was profiled by 16S rRNA amplicon sequencing of feces from SPF WT mice fed BP or PP diets for 1 week in the absence of colitis. Principal coordinates analysis demonstrated distinct clustering of mice according to dietary protein source, which explained approximately 65% of variation in the data (PERMANOVA test coefficient of determination, R^2^ = 0.43, p=0.005) (**Fig 3A**). Although profiles before feeding either experimental diet were very similar, 7 days of feeding BP or PP diet significantly altered the taxonomic abundance of multiple bacteria (**Fig 3B; Fig. S2**). Significant expansion of *Akkermansia muciniphila* occurred in the BP diet group, while *Turicibacter sanguinis* increased in the PP diet group, along with preservation of *Lactobacillus johnsonii* (**Fig. 3B-C**). *A. muciniphila* can impair gut barrier function through mucus digestion^12, 17, 21, 28–30^, therefore, we examined its colonic luminal spatial distribution by fluorescence *in situ* hybridization (FISH). This revealed high abundance of *A. muciniphila* adjacent to the mucus layer of BP-fed mice while no signal was observed in PP-fed mice (**Fig. 3D**). Colonic surface mucus thickness showed a corresponding ~50% reduction in BP *vs*. PP fed mice (**Fig. 3E**). Additionally, surface mucus quality was reduced in BP-*vs*. PP-fed mice with no obvious difference in goblet cell mucus staining (**Fig. 3F; Fig. S3**). To evaluate whether BP-driven expansion of *A. muciniphila* was responsible, in part, for promoting colitis, GF *Il10*^−/−^ mice were selectively colonized with *A. muciniphila* alone, a consortium of IBD-associated pathobionts including *Escherichia coli*, *Enterococcus faecalis*, and *Ruminococcus gnavus* (EER), or *A. muciniphila* plus EER and fed BP diet. Although *A. muciniphila* mono-association did not induce colitis, it significantly potentiated EER pathobiont-induced colitis severity (**Fig 3G-H**). Appropriate strain colonization was confirmed by anaerobic plating of feces from colonized mice followed by 16S colony PCR and Sanger sequencing of representative colony morphologies (data not shown). These data suggest that BP exacerbates colitis, in part, by promoting expansion of mucus-digesting *A. muciniphila* that potentiates the pro-inflammatory effects of IBD-associated pathobionts.

**Figure 3.**
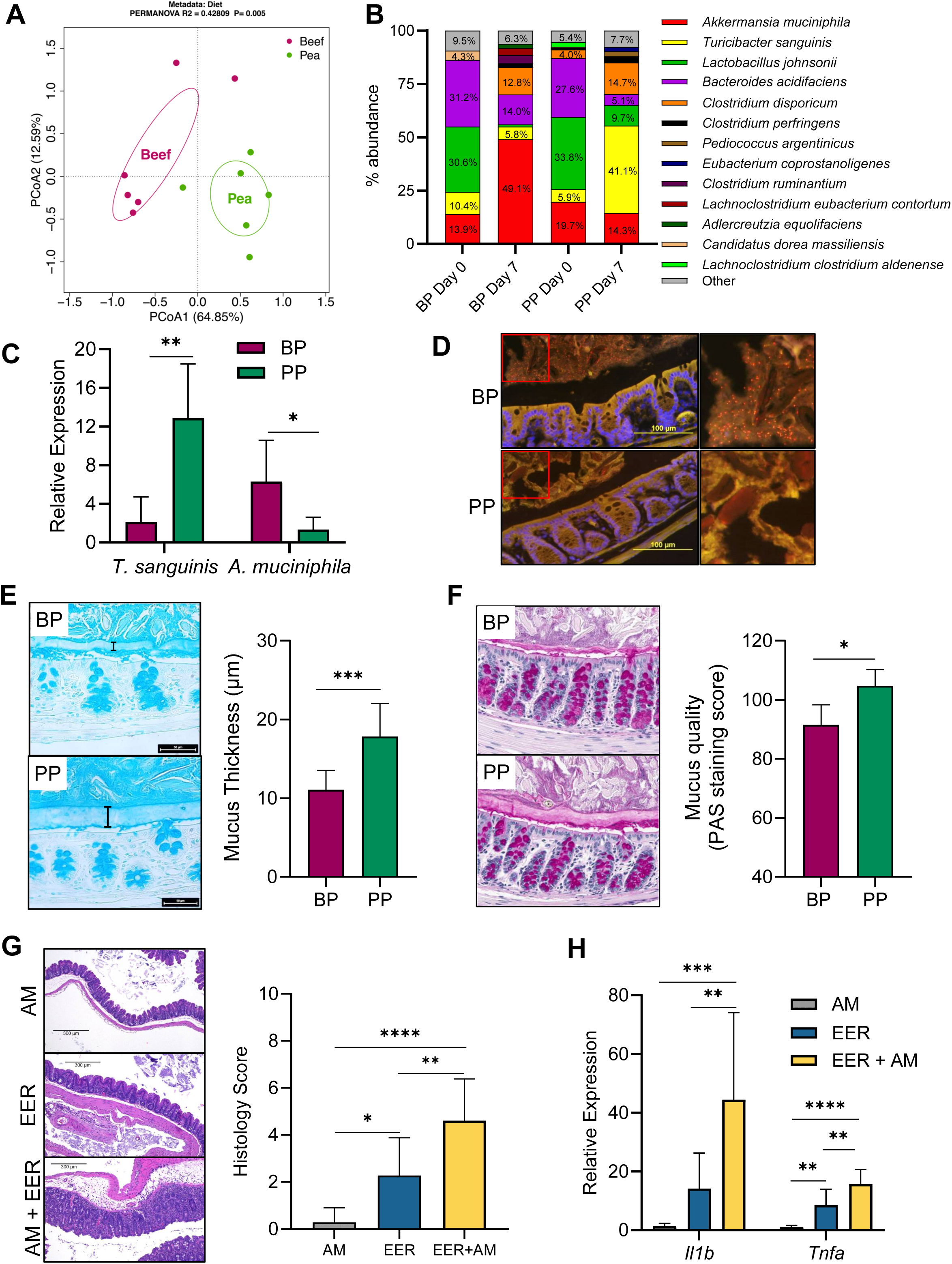
Beef protein consumption promotes *A. muciniphila* expansion and gut barrier defects. A-C, C57BL/6J mice were fed either beef protein (BP) or pea protein (PP)-containing diets for 7 days then fecal samples were subjected to 16S rRNA sequencing. Results are displayed as principal coordinate analysis on day 7 (A) and as percent abundance of each species on days 0 and 7 (B) (those species <1% in overall abundance are labeled as ‘Other’). qRT-PCR was performed on fecal DNA to measure relative abundance of major populations found in either diet group (C). D-F, C57BL/6J wild-type mice were fed BP or PP diets for 7 days then colons were harvested and fixed to preserve the mucus layer. Fluorescence *in situ* hybridization was performed to identify *A. muciniphila* (indicated by orange structures) (D), Alcian blue staining was carried out to measure mucus thickness (representative images shown on left side and quantification of staining shown on right side; bracket indicates mucus layer on images) (E), and periodic acid Schiff staining was performed to quantify mucus quality (representative images shown on left side and quantification of staining shown on right side) (F). n = 6 per group. G-H, germ-free *Il10^−/−^* mice were inoculated with *A. muciniphila*, a defined consortium of *Escherichia coli*, *Enterococcus faecalis*, and *Ruminococcus gnavus* alone or in combination with *A. muciniphila* and fed BP diet. Representative histologic images of ceca are shown (left side) and total histology score was assessed (right side) (G) and relative expression of inflammatory cytokine genes was measured (H) in the ceca of mice on day 14. n = 7-9 per group, data representative of 2 independent experiments. *P<0.05; **P<0.01; ***P<0.001; ****P<0.0001. Bar plots show mean and standard deviation.

Because bacteria have an important role in mediating BA metabolism and composition and there was higher abundance of bile salt hydrolase (BSH)-carrying^31–34^ bacteria (*T sanguinis*, *L johnsonii*) in PP-fed mice, we next measured the relative levels of common BAs in feces from WT SPF mice fed BP or PP diets for 7 days in the absence of colitis. This analysis showed that most BAs in BP-fed mice were primary and conjugated while most BAs in PP fed mice were secondary and unconjugated (**Fig. 4A**). Multiple taurine conjugated primary BAs, including taurocholic acid (TCA), were significantly higher in BP-fed mice while several unconjugated secondary BAs, including lithocholic acid (LCA) and deoxycholic acid (DCA), were elevated in PP-fed mice (**Fig. 4B**). These patterns were not observed in the livers or ileal content of mice fed BP *vs*. PP diets (data not shown), suggesting that fecal BA alterations are likely mediated by colon-specific microbial differences. To connect the respective BA profiles of mice fed PP or BP diets with differential bacterial abundance, we performed an *in vitro* bile acid deconjugation assay comparing select bacterial species enriched by either diet. This revealed that *L. johnsonii* (higher abundance in PP-fed mice) deconjugated TCA to cholic acid (CA) 100-fold more than *A. muciniphila* (increased in BP-fed mice) (**Fig. 4C**). To evaluate whether the primary CBAs that were increased in feces of BP diet-fed mice directly exacerbate colitis, PP-fed mice were administered TCA by oral gavage daily for 7 days and exposed to DSS. DAI, colon length and histologic score showed that TCA worsened colitis in PP-fed mice (**Fig 4D-F**). Subsequent MALDI-MSI based imaging of the colon of mice administered TCA in the absence of colitis revealed TCA accumulation in the colonic epithelium (**Fig. S4**). Collectively, these data suggest that mice consuming a BP-containing diet have lower abundance of BA-deconjugating bacteria, resulting in higher levels of colitis-promoting primary CBAs.

**Figure 4.**
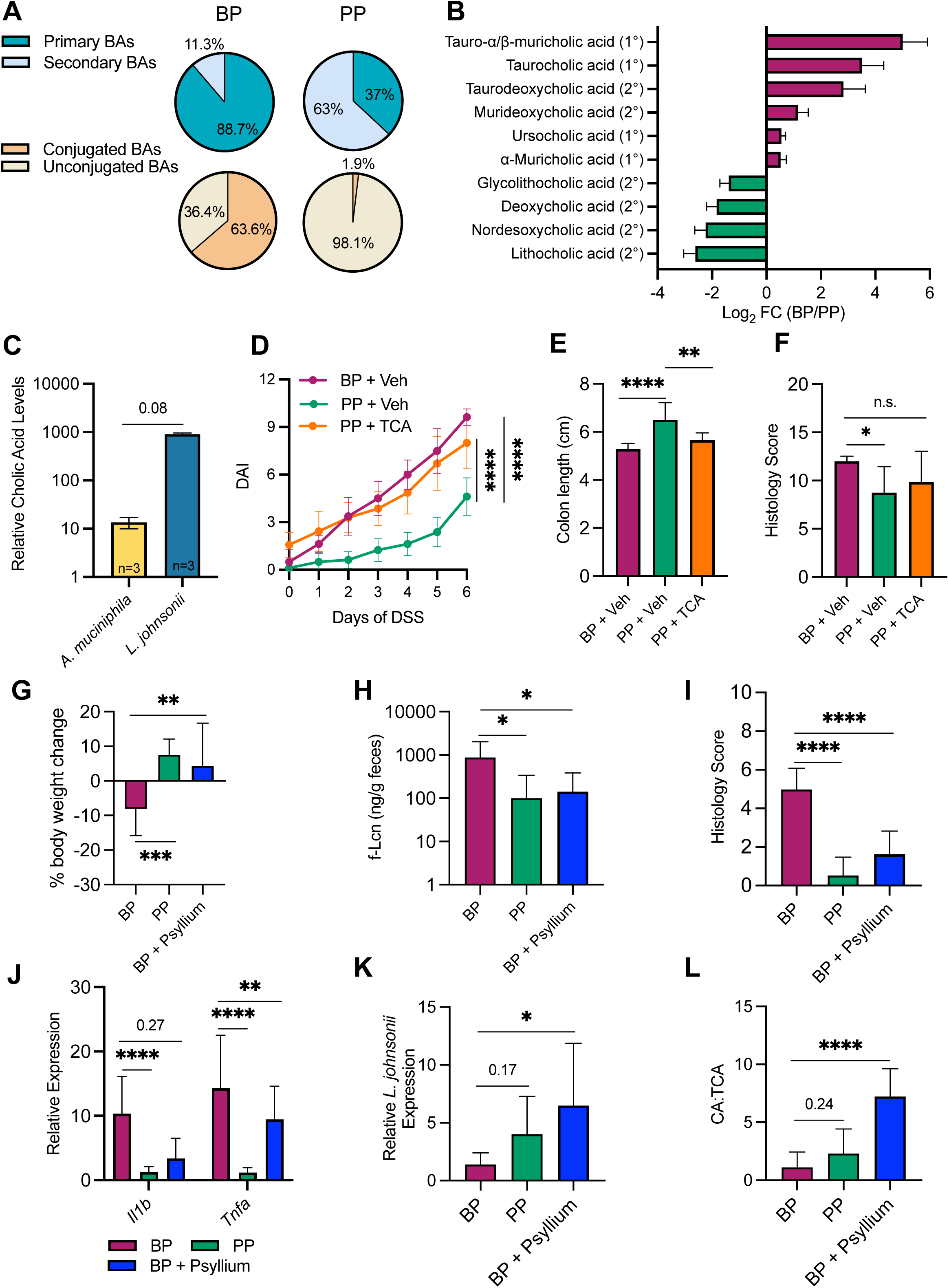
Bacteria-mediated bile acid changes facilitate colitis severity. A-B, WT C57BL/6J mice were fed diets containing either beef protein (BP) or pea protein (PP)-containing diets for 7 days and the relative abundance of bile acids was measured in feces. The average percent of primary *vs*. secondary or conjugated *vs*. unconjugated bile acids in each diet group is shown (A) and those BAs that displayed significantly different abundance (FDR<0.1) are shown as log2 fold-change in PP-fed mice relative to BP-fed mice (B). C, taurocholic acid (TCA) (1mM) was administered to culture medium of *L. johnsonii* and *A. muciniphila* and the levels of cholic acid were measured in the supernatants after 24 hours and reported as relative to either species cultured in the absence of TCA. D-F, C57BL/6J WT mice were fed either BP or PP diets and given daily oral gavage of PBS or fed PP and gavaged daily with TCA (0.35mg/kg) for 7 days, then administered 1% DSS in drinking water for 7 days, while continuing oral administration. Disease Activity Index (DAI) was measured over 7 days (D), colon length was measured on day 7 (E) and histologic inflammation of colons was assessed on day 7 (F). G-J, germ-free *Il10^−/−^* mice were inoculated with mouse-adapted pooled human IBD patient fecal microbiota (IMM-HM2) then fed BP or PP diets containing cellulose or BP diet containing psyllium (BP + psyllium) for 14 days. Weight changes relative to day 0 were calculated (G), fecal lipocalin-2 (f-Lcn) was measured (H), total histology score was assessed (I), and relative gene expression of inflammatory cytokines in colons was measured (J), on day 14. n = 10-11 mice per group. K-L, WT specific pathogen-free mice were fed the diets described in panel G for 7 days and feces were collected to measure the relative abundance of *L. johnsonii* by qRT-PCR (K) and the ratio of cholic acid to taurocholic acid (CA:TCA) in fecal samples (L). n = 10-11 mice per group. *P<0.05; **P<0.01; ***P<0.001; ****P<0.0001. Bar plots show mean and standard deviation.

Microbe-accessible dietary fiber supports populations of gut health-promoting commensals, including BA-metabolizing bacteria^35^. Therefore, we reasoned that maintaining these bacterial populations by adding soluble dietary fiber would attenuate BP-driven colitis severity. To test this concept, ex-GF *Il10*^−/−^ mice inoculated with MA human IBD microbiota were fed PP diet, BP diet, or BP diet in which cellulose fiber was replaced with psyllium (BP-psyllium diet) and monitored for colitis severity (**Table S1**). Compared to BP diet, the BP-psyllium diet significantly decreased colitis severity, as determined by body weight, fecal lipocalin, histologic analysis and cytokine gene expression (**Fig. 4G-J**). Similar protection by psyllium was observed in the DSS-induced colitis model (**Fig. S5**). Analysis of BA-metabolizing bacteria and BA profiles in WT C57BL/6J mice fed each of these three diets in the absence of colitis showed that psyllium supplementation significantly increased the relative abundance of *L. johnsonii* and the CA:TCA ratio in BP-fed mice (**Fig. 4K-L**). Taken together, these data support a model in which BP-exacerbated colitis, which is driven by high abundance of primary CBAs, can be attenuated through psyllium fiber administration, restoring populations of BSH-carrying bacteria that convert primary CBAs to their deconjugated form.

Plant-based diets are believed to benefit IBD patients, while animal-based diets may be detrimental. Both retrospective and prospective studies suggest that consumption of red meat is associated with IBD incidence and disease activity. For example, The E3N Prospective Cohort Study revealed that high animal protein consumption significantly increased IBD incidence^36^ and multiple studies showed that meat intake increased the risk of disease flare in patients with ulcerative colitis (UC)^5, 37, 38^. The concept that plant-based diets are beneficial largely stems from studies demonstrating that microbe-fermentable carbohydrates from plants reduce experimental colitis^2, 39^. In contrast, diets rich in animal-derived products (especially red meat) are thought to promote intestinal inflammation through heme, sulfur and saturated fat content^2, 39^. Although these components of plant and animal-based diets impact gut health, the role of protein content has not been widely considered. Our study, which selectively examined protein content by feeding mice identical diets except for the type of protein isolate, suggests that the source of dietary protein content differentially impacts colitis severity. Other studies investigating the impact of protein source on experimental colitis, including from beef and pea, are largely consistent with the data shown in the current study, although this previous work used diets containing whole food sources rather than purified protein isolate^14, 15, 40, 41^. Our findings support a role for the dietary protein-altered microbiota and BAs in mediating colitis severity, however, the specific mechanism(s) by which protein type drives these effects remains unclear. It is plausible that differences in amino acid composition or digestion/absorption of diverse dietary protein sources are contributing factors. Therefore, determining which amino acids or what properties of each protein type mediate intestinal inflammation will be important for future studies.

Bacteria, BAs and their interplay are key mediators of IBD pathogenesis^1, 2^. Higher levels of conjugated and primary BAs are found in feces of patients with active IBD, compared to unconjugated and secondary BAs in healthy individuals^23–25^. Our work suggests that a shift to a predominance of conjugated and primary BAs, is in part, responsible for greater colitis severity when feeding BP to mice. We found that taurine conjugated and primary BAs are higher in BP-fed mice than PP-fed mice, and showed that direct administration of TCA, which was highly increased in the BP-fed group, worsened colitis in PP-fed mice to levels observed in mice given BP diet alone. Although the exact mechanism(s) by which altered BAs enhance disease is unclear, the detergent-like properties of BAs could disrupt gut barrier function by impairing epithelial cell function and/or epithelial-protective mucus. This concept is supported by our data showing the ability of exogenously administered TCA to localize to the colonic epithelium.

Interestingly, our prior work showed that a high fructose diet exacerbated murine colitis with a similar increase in CBAs as observed in BP fed mice, with rectal administration of CBAs worsening colitis and thinning colonic mucus in mice fed a control diet^17^. Integrating our study focused on dietary protein with prior studies of the role of dietary fat and carbohydrates in experimental colitis suggests that altered BAs may be a common pathway for diet-mediated gut inflammation^17, 22^. Although our work focused on the detrimental impact of conjugated primary bile acids enriched in BP-fed mice, the profile of unconjugated, secondary bile acids enriched in feces from PP fed mice might itself be protective. For example, LCA and DCA, both enriched in PP-fed mice, are strong ligands of the PXR and TGR5 receptors that, when activated, induce anti-inflammatory effects^24, 42–44^. Therefore, the role of unconjugated secondary bile acids as protective metabolites in PP-fed mice is an important future direction.

Bile acids have a bidirectional relationship with gut bacteria, whereby bacterial enzymes metabolize BAs, which in turn, alter bacterial physiology and microbiota composition^42^. PP feeding expanded bacterial species that efficiently deconjugate BAs, including *L johnsonii* and *T sanguinis* (as shown by data from the deconjugation assay performed in the current study and previous work^32^), while BP feeding expanded *A muciniphila,* a poor deconjugator that also degrades mucus and potentiates pathobiont-driven inflammation^45^. These dietary protein-induced bacterial shifts likely caused the observed BA profiles, however, we cannot exclude the possibility that altered BAs directly shifted microbiota composition, as previously demonstrated^46–49^. Further evidence that the ratio of conjugated to unconjugated BAs are important for mediating colitis stems from our findings that psyllium supplementation protected against BP-mediated colitis severity, while maintaining favorable BA profiles and populations of BA-deconjugating bacteria. However, psyllium fiber likely protects against inflammation by multiple mechanisms, including SCFA production and prevention of bacterial mucus degradation^12, 50^.

Our study reaffirms the concept that IBD occurs when multiple detrimental factors intersect to initiate and perpetuate gut inflammation^51^. These IBD determinants – genetically dysregulated host inflammatory and epithelial barrier responses, abnormal resident microbes, and environmental triggers – can exist in isolation or combination without causing disease, but together drive severe IBD. For example, GF colitis susceptible *Il10^−/−^* mice fed BP diet (genetic susceptibility + environmental trigger) did not develop inflammation, nor did SPF WT mice fed BP-diet (resident microbes + environmental trigger). However, we demonstrated that when all three IBD-driving factors – host, microbe, and environment – intersect, severe and progressive intestinal inflammation occurs. The poor long-term success of current human IBD therapies may reflect a failure to reconcile the multiple determinants of IBD. For example, addressing just one factor, such as suppression of host inflammation, leaves the patient vulnerable to recurrent disease upon encountering environmental triggers. We suggest that durable long-term remission requires simultaneously addressing all IBD determinants, and as such, investigations into environmental drivers of IBD, especially diet, is required to guide clinical approaches to maintain sustained remission.

## Methods

### Murine colitis models

To induce chemical colitis, 8-week-old male C57BL/6J mice (The Jackson Laboratory) were administered 1% DSS (Sigma) in drinking water for 6–7 days, as indicated. Colitis severity was assessed by measuring changes in body weight, as well as severity of rectal bleeding and diarrhea, as previously described^17, 52^. Disease Activity Index (DAI) was calculated by combining severity of body weight loss, diarrhea and bleeding, as previously described^17, 52^.

T cell-mediated colitis was induced by inoculation of mouse-adapted human IBD patient fecal microbiota (previously developed and characterized^27^) to GF *Il10^−/−^*mice on a 129S6/SvEv background (purchased from the National Gnotobiotic Rodent Resource Center (NGRRC) at the University of North Carolina), as previously described^27^. Mouse-adapted (MA) human IBD patient microbiota was previously generated from de-identified human IBD patient stool samples collected under an Institutional Review Board approved protocol; no primary human samples were used in this study^27^. Briefly, human fecal materials from pooled cohorts of human donors with active IBD were passaged through GF 129S6/SvEv *Il10^−/−^* mice to generate standardized aliquots of fecal slurry^27^. MA IMM-g2 microbiota was derived from pooled feces of 2 CD and 1 UC patient and MA IMM-HM2 microbiota was derived from pooled feces of 3 CD patients^27^.

Standardized 100mg/ml aliquots of mouse-adapted human IBD microbiota were anaerobically thawed, diluted with pre-reduced PBS, and administered by oral gavage (2mg) to recipient GF *Il10^−/−^* mice^27^. All fecal transplant experiments were performed with sterile gnotobiotic cage technique in BSL-2 isolation cubicles with HEPA-filtered air^53^. Prior to colonization, GF mice were fed Purina Advanced Protocol Select Rodent 50 IF/6F Auto Diet. Defined diets were started at the time of fecal microbiota transplant.

At the end of the experimental periods, mice were humanely euthanized using CO_2._ Excised colons were measured then flushed with ice-cold phosphate-buffered saline (PBS). Tissue was either snap-frozen for biochemical analysis or fixed in 4% paraformaldehyde or 10% phosphate buffered formalin for 4-24 hours, followed by paraffin embedding and H&E staining for histological analysis. All animal studies were approved by the Institutional Animal Care and Use Committee at Stony Brook University and the University of North Carolina at Chapel Hill.

### Diet-polarized fecal microbiota transplant (FMT)

Feces were collected from WT C57BL/6J SPF mice fed BP or PP diets for 2 weeks, anaerobically homogenized, and diluted in pre-reduced lysogeny broth (LB) with 20% glycerol to generate fecal slurries (n=5-6 donor mice per diet). Each donor FMT was administered to 1-2 recipient mice GF C57BL/6J mice via oral gavage on days 0, 3, 5, 7, and 9 of the experiment. Mice were administered 2% DSS from days 3-10 of the experiment and monitored for signs of colitis.

### Inoculation of defined consortia

*Escherichia coli* LF82^54^, *Enterococcus faecalis* OG1RF^55^, and *Ruminococcus gnavus* ATCC 29149^55^ (EER) were grown under anaerobic conditions in brain-heart infusion medium supplemented with 5g/L yeast extract, 0.5g/L L-cysteine, and 5mg/L hemin (LYH-BHI medium). *A. muciniphila* ATCC BAA-835 (AM) was grown under anaerobic conditions in LYH-BHI medium supplemented with 5g/L porcine gastric mucin (LYH-BHI + PGM). Pure cultures of each strain were grown anaerobically to confluence then equally mixed to generate EER or EER + AM consortia. GF 129S6/SvEv *Il10^−/−^*mice were colonized with freshly cultured consortia by oral gavage and given BP diet from the time of colonization. Feces were collected at necropsy, serial diluted and plated under anaerobic conditions on LYH-BHI + PGM to confirm colonization with consortia members. Representative colonies of each strain identified by colony morphology were picked from plates, 16S PCR amplified, and Sanger sequenced (Eton Bioscience) to confirm colonization of appropriate strains in recipient mice.

### Histopathologic scoring

Colons from mice given DSS were flushed, formalin fixed, and paraffin embedded followed by sectioning and staining with hematoxylin and eosin. Scoring of colitis/colonic inflammation was performed based on evaluation of mucosal damage by acute inflammation, by crypt abscess, mucosal architectural distortion and proportion of the involvement of colon as previously described^56^. Briefly, epithelial damage by acute inflammation was graded into 4 categories (0-3; 0=no inflammation. 1=mild, 2=moderate and 3=severe inflammation with ulceration). Mucosal damage with crypt abscess graded into 2 categories (0=absent and 1=present). Extent of acute inflammation was graded into 4 categories (0-3; 0=no inflammation, 1=mucosal, 2=submucosal and 3=transmural). Mucosal architectural distortion was graded into 4 categories which includes the percentage of length of colon involvement (0-3; 0=normal architecture, 1= mild/focal or 10%, 2=moderate/20-30%, and 3=severe/ >40% of the entire length). Proportion of the total involvement was evaluated by the percentage of length of colon involved by colitis into 4 categories (0-3; 0=no colitis, 1=1-10% of the total length, 2=20-30% and 3=>40% of the total evaluated length of colon). All histologic scoring was performed in a blinded fashion by a board-certified gastrointestinal pathologist (S.A.).

For *Il10^−/−^* experimental colitis studies, histologic inflammation was quantified in ileal, cecal and colonic (proximal, distal, rectal) tissue segments using a well-validated rubric by blinded histopathology scoring on a scale of 0-4 for each segment, as previously described^27, 56,57^. Total histology score was calculated by summation of all 5 tissue segment scores for a scale of 0-20.

### Fecal lipocalin-2 quantification

Fecal samples were homogenized in PBS with 0.1% Tween 20, incubated at 4°C for 12 hours, then centrifuged to yield clear supernatant for lipocalin-2 ELISA, performed according to the manufacturer’s instructions (DY1857, R&D Systems)^58^.

### Measurements of mucus thickness and quality

Segments of harvested colons from mice containing a fecal pellet were preserved in fresh Carnoy’s fixative solution (methanol:chloroform:glacial acetic acid, 60:30:10) then sequentially washed in methanol, ethanol and xylenes, as previously described^17^. Tissues were then paraffin-embedded and sectioned (5 μm), followed by staining with Alcian Blue, and mucus thickness was measured in 20-50 regions of each colon in a blinded manner using ImageJ software (National Institutes of Health, Bethesda, MD), as previously described^17^.

To determine differences in mucus quality on the surface of the epithelium and within goblet cells, Periodic acid Schiff (PAS) staining was performed. Stained slides were scanned using an Aperio Image Scope (ScanScope, Aperio, CA) and used to count the PAS-positive pixels by the Aperio Color Deconvolution algorithm, as previously described^17^.

### qRT-PCR

For measurements of mammalian gene expression, RNA was isolated from tissues using QIAzol Lysis Reagent (Qiagen) following the manufacturer’s protocol. RNA concentration was measured using a NanoDrop ND-1000 spectrophotometer. cDNA was synthesized using 2000 ng total RNA and reverse transcriptase qScript SuperMix (Quanta). Quantitative real-time PCR was performed using specific primers on a QuantStudio 7 Real-Time PCR System (Applied Biosystems) using PowerTrack SYBR Green PCR Master Mix and oligo (dT) primers according to the user’s manual. *Actb* expression was used as internal control for gene expression. The ΔΔCT method was used to calculate relative fold expression. Primer sequences are shown in **Table S2**.

For bacterial abundance and bacterial gene expression analysis, DNA was extracted from fecal samples using the QIAamp DNA Fast Stool Mini Kit (Qiagen) per the manufacturer’s instructions. For each sample, 10 ng of DNA was used for qRT-PCR using PowerTrack SYBR green PCR master mix on a QuantStudio 7 PCR Machine. Bacterial primers sequences are shown in **Table S2**. Relative abundance was calculated by the ΔΔCT method using universal 16S primers as control.

### Bile acid analysis

Fecal and ileal samples were dried in a vacuum centrifuge then homogenized in 500 µL of cold 80% methanol. Liver samples were homogenized using the same method but without prior drying. Bacterial supernatants were mixed with cold 80% methanol in a 1:9 ratio. Following homogenization or mixing, samples were centrifuged at 14,000 rcf for 20 minutes at 4°C and the supernatant was collected and dried down using a vacuum centrifuge then stored at −80° C prior to analysis. Bile Acid analysis was performed as previously described on a Vanquish UHPLC system coupled to a Q Exactive Orbitrap mass spectrometer (Thermo Scientific)^17^. Relative quantitation was normalized to dry weight of fecal/ileal samples or protein concentration of livers. Analysis was performed at the Proteomics & Metabolomics Core Facility of Weill Cornell Medicine.

### MALDI Mass Spectrometry Imaging

Tissue preparation and imaging was performed following a previously published method^59^. Briefly, mouse colons were processed into Swiss-rolls, embedded in 5% carboxymethylcellulose, and stored at −20°C. Serial frozen sectioning was performed for paired MALDI imaging and standard H&E staining. MALDI imaging was conducted on a rapifleX mass spectrometer (Bruker Daltonics) within metabolite mass range in negative ion detect mode at 50 µm spatial resolution. The TCA deprotonated ion with a mass-to-charge ratio of 514.284 m/z was detected, and the average intensity of the mass spectrometry signal for TCA determined using SCiLS Lab software (Bruker Daltronics).

### 16S rRNA Analysis

Frozen fecal samples were shipped to Molecular Research (Shallowater, TX) for 16S rRNA profiling, performed as previously described^17^. DNA was extracted with the Powersoil DNA Kit (Qiagen). The 16S rRNA gene V4 variable region was targeted for PCR amplification, followed by Illumina HiSeq short-read sequencing. Sequencing outputs were processed and taxonomically classified as previously described^17^. Operational taxonomic units were defined by clustering at 97% similarity (3% divergence) and classified by BLASTn against an RDPII/NCBI derived database. Statistical analysis and visualization of principal coordinates analysis (PCoA) plots were performed with the Shiny application *Plotmicrobiome* (Sun et al. GitHub https://github.com/ssun6/plotmicrobiome).

### Fluorescence in situ hybridization (FISH)

Fluorescence in situ hybridization (FISH) was performed as previously described^17, 60^ on 4μm histologic sections of Carnoy’s-fixed paraffin-embedded colon tissue using an *A. muciniphila*-specific FISH probe (S-SMUC-1437-a-A-20: Cy-3-50-CCTTGCGGTTGGCTTCAGAT-30)^17, 60^. Slides spotted with multiple control bacteria (*E. coli, P vulgaris, K pneumoniae, S equi, S bovis*) not-reactive to the A.M. FISH probe were used to confirm probe specificity^17, 60^.

### In vitro bacterial culture and treatment

*L. johnsonii* BAA-3147 and *A. muciniphila* BAA-835 from frozen stocks were cultured overnight in MRS or LYH-BHI + 0.5% mucin, respectively, the day before the experiments. *A. muciniphila* BAA-835 was cultured in an anaerobic chamber (Coy Laboratory) with mixed gas (90% N2, 5% CO2 and 5% H2; 37°C), and *L. johnsonii* BAA-3147 was cultured under aerobic conditions (37°C with shaking at 150 rpm). The overnight cultures were diluted with fresh media with or without 1 mM taurocholate to *A. muciniphilla* BAA-835 (once at OD600 of 1-1.1) and to *L. johnsonii* (once at OD600 of 0.7-0.8). The diluted cultures were immediately distributed into two sets of tubes for 0 and 24 h time points, each in triplicate. The time 0 h samples were transferred to ice immediately to prevent bacterial activity on taurocholate. Optical densities were measured at each time point for normalization. The cultures were then centrifuged at 3,000 rpm for 20 min at 4°C and the supernatants were filtered through a 0.2 µm syringe filter and kept at −80°C until BA measurements.

### Statistical analysis

For the comparison of DAI across different mice groups, linear mixed-effects models for longitudinal data analysis were used. Model assumption was confirmed using residual diagnosis. Time (in days) was treated as a continuous variable to assess the trend in DAI. An interaction term between diet group and time was used to compare the trends across groups. The covariance structure used to model the correlated longitudinal measurements from the same mice was unstructured (UN), selected based on Akaike Information Criterion (AIC) from a set of possible structures. For the comparison of numeric values observed or calculated at the end of experiments, Wilcoxon rank-sum tests or simple linear regression models were used.

These analyses were applied to the following variables: % body weight change, histology score, gene expression, colon length, bacterial abundance, mucus thickness, mucus quality, and bile acid ratios. A P-value less than 0.05 was considered statistically significant and analyses were performed using SAS 9.4 (SAS Institute Inc., Cary, NC).

#### Statistical analysis of BA levels

The BA dataset was filtered to remove BA with >50% missing values. Normalization of raw metabolite levels was performed by probabilistic quotient normalization followed by log2-transformation and imputation of missing values by k-nearest neighbor (KNN). BA differences were quantified by log2 fold change from the PP diet group. Linear models with diet as the independent variable were used to calculate statistical significance. The Benjamini-Hochberg method with an FDR cut-off of 0.2 was used for multiple hypothesis correction. Analyses were performed with the maplet package (version 1.2.1) in R ^61^.

## Acknowledgements

The authors thank members of the Montrose and Sartor laboratories for suggestions, helpful comments, and technical support. We thank Josh Frost, manager of the National Gnotobiotic Rodent Resource Center, and the Center staff members for gnotobiotic husbandry and mice. We acknowledge the Biological Mass Spectrometry Shared Resource at the Stony Brook Cancer Center for expert assistance with the MALDI-MS imaging analysis. We acknowledge the biostatistical consultation and support provided by the Biostatistical Consulting Core at School of Medicine, Stony Brook University.

## Supplementary Figure Legends

**Figure S1.**
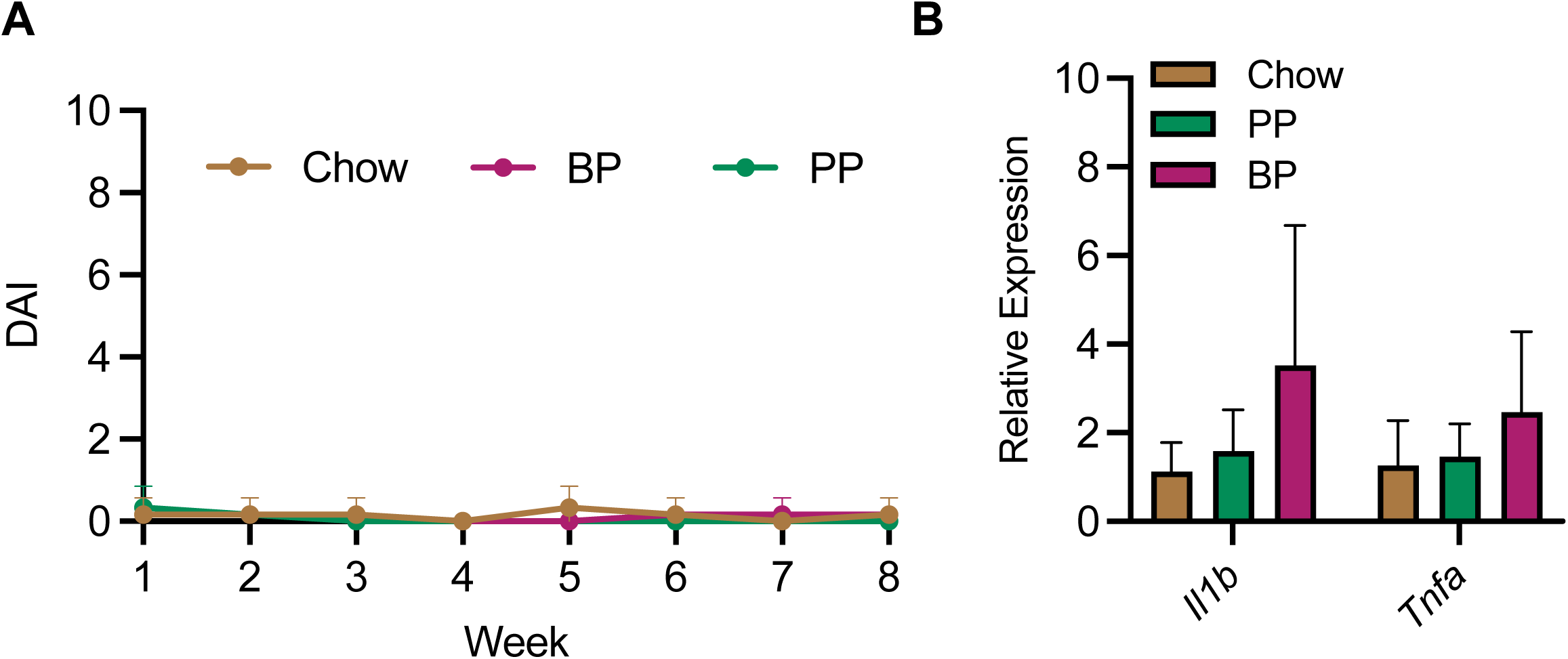
Long-term beef protein feeding in specific pathogen free wild-type mice does not induce signs of colitis. Mice were fed chow, or purified diet containing beef protein (BP) or pea protein (PP) isolate for 8 weeks. Disease activity index (DAI) was measured weekly (A) and relative expression of *Tnfa* and *Il1b* was measured in colons at the end of the study (B). n = 6 mice per group. Bar plots show mean and standard deviation.

**Figure S2.**
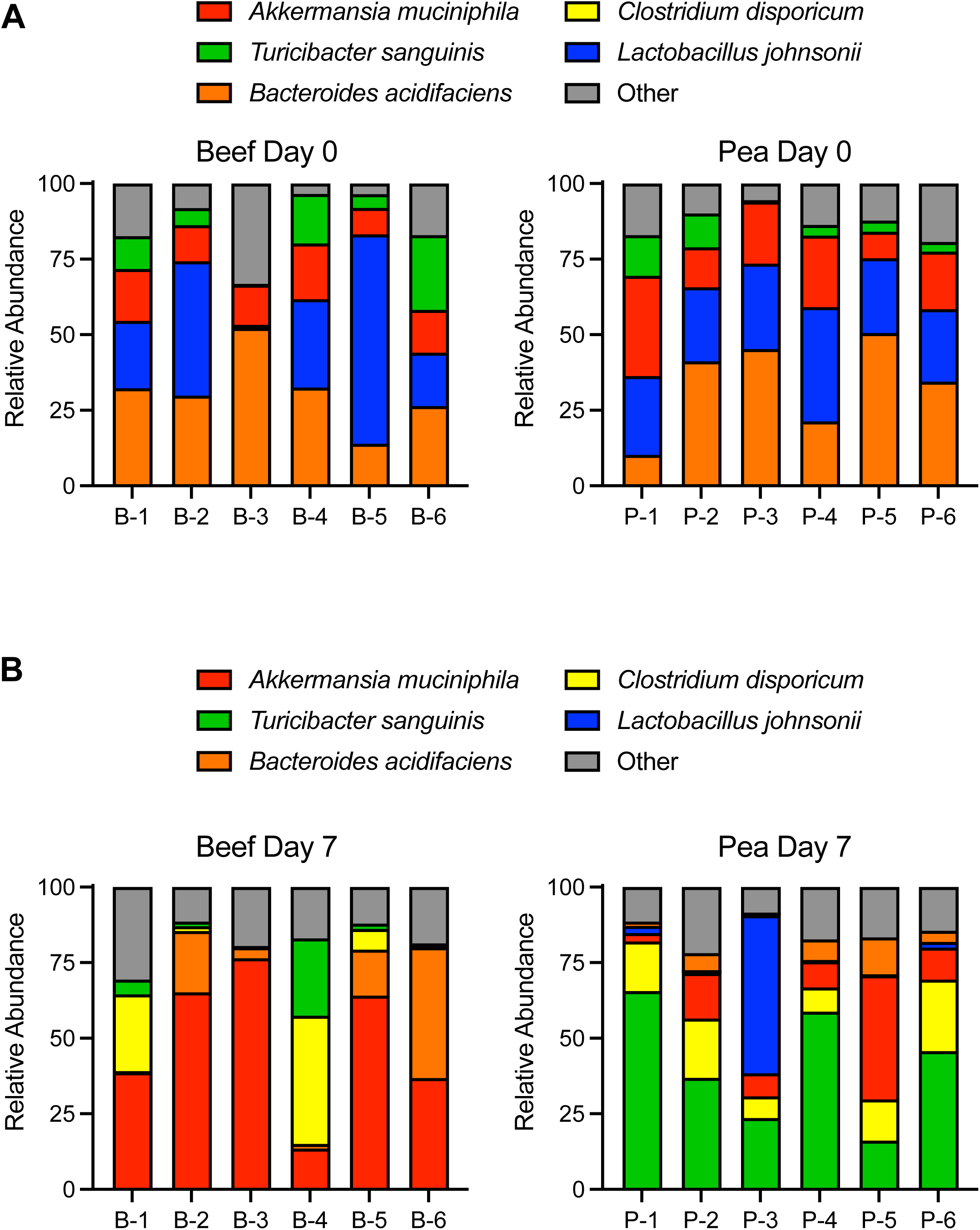
Relative bacterial species abundance of individual mice fed beef or pea protein-containing diets. Species abundance in feces from individual mice is reported prior to (A) and 7 days following (B) administration of diets containing beef protein (BP) or pea protein (PP) isolate. Those species appearing in abundance <1% in all conditions are labeled as ‘Other’.

**Figure S3.**
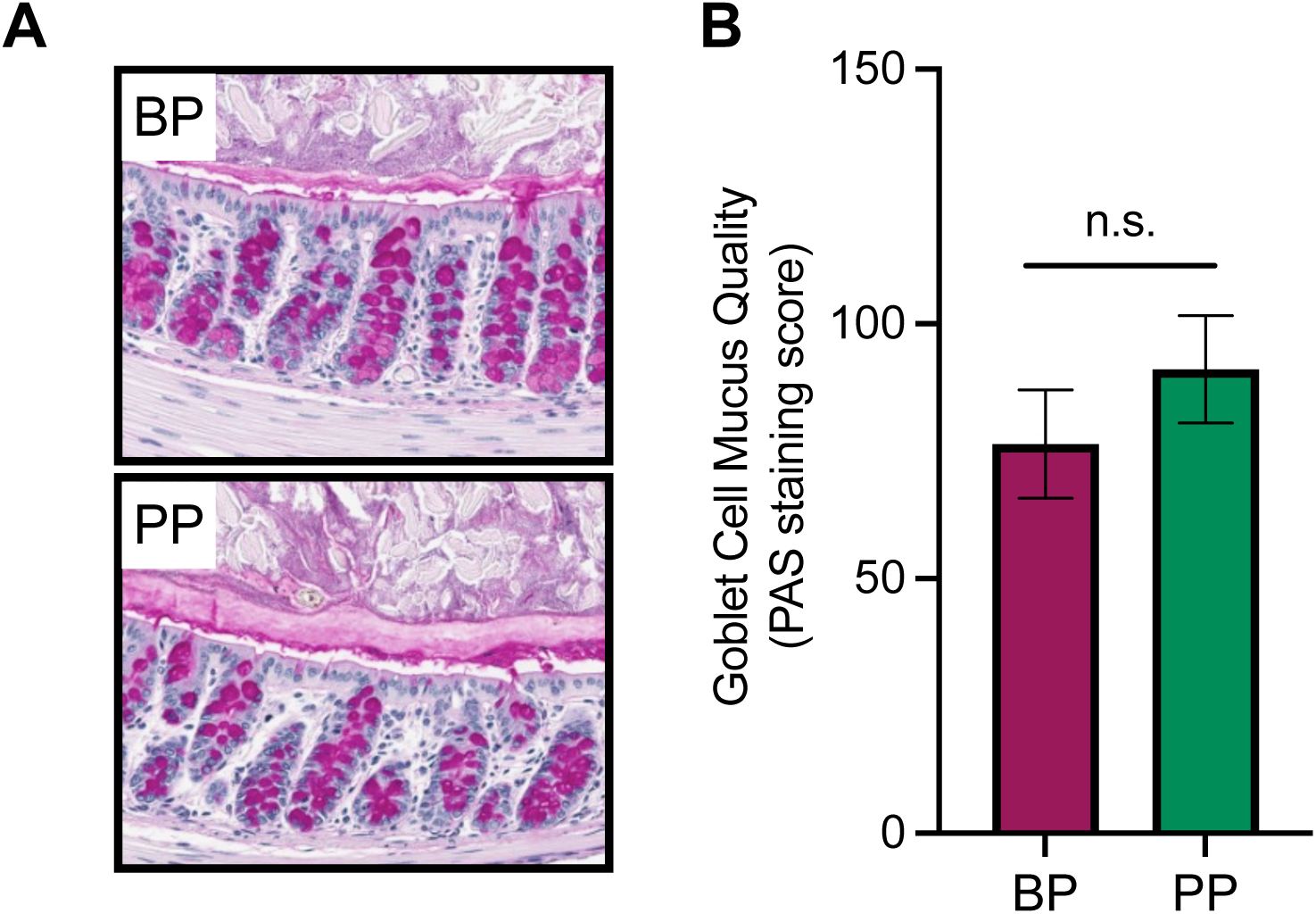
Dietary protein source does not affect goblet cell mucus quality. C57BL/6J mice were fed either beef protein (BP) or pea protein (PP)-containing diets for 7 days then periodic acid Schiff staining was performed to quantify mucus quality within goblet cells. Representative images (A) and quantification of staining (B) are shown. Bar plots show mean and standard deviation.

**Figure S4.**
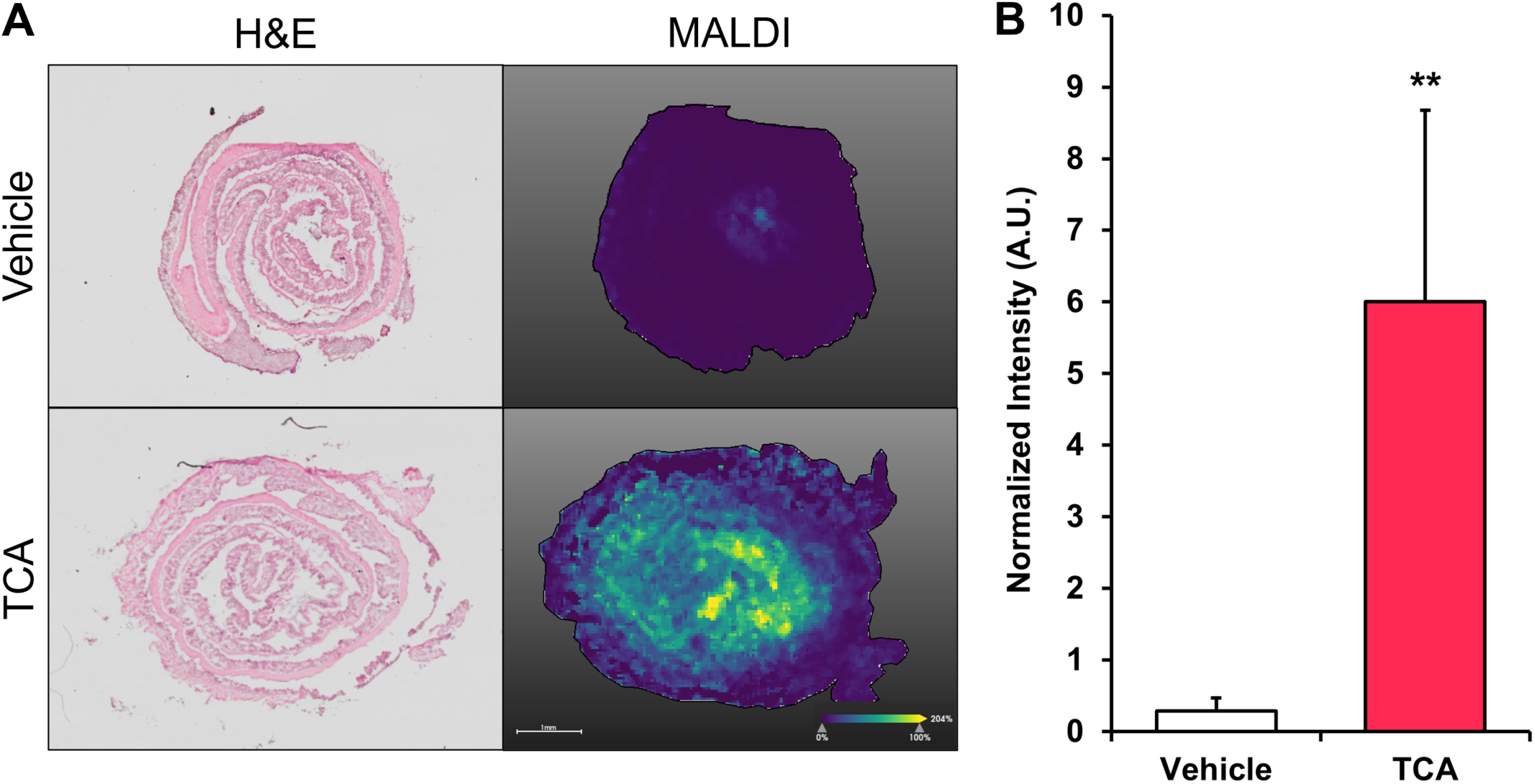
Taurocholic acid localizes to the colonic epithelium following oral administration to mice. C57BL/6J WT mice were administered daily oral gavage of PBS or taurocholic acid (TCA) (0.35mg/kg) for 7 days and colon tissue sections were subjected to MALDI-MSI to determine localization (A) and relative abundance (B) of TCA in each group. TCA group is normalized to vehicle group in panel B. n = 4 samples per group. **P<0.01. Bar plots show mean and standard deviation.

**Figure S5.**
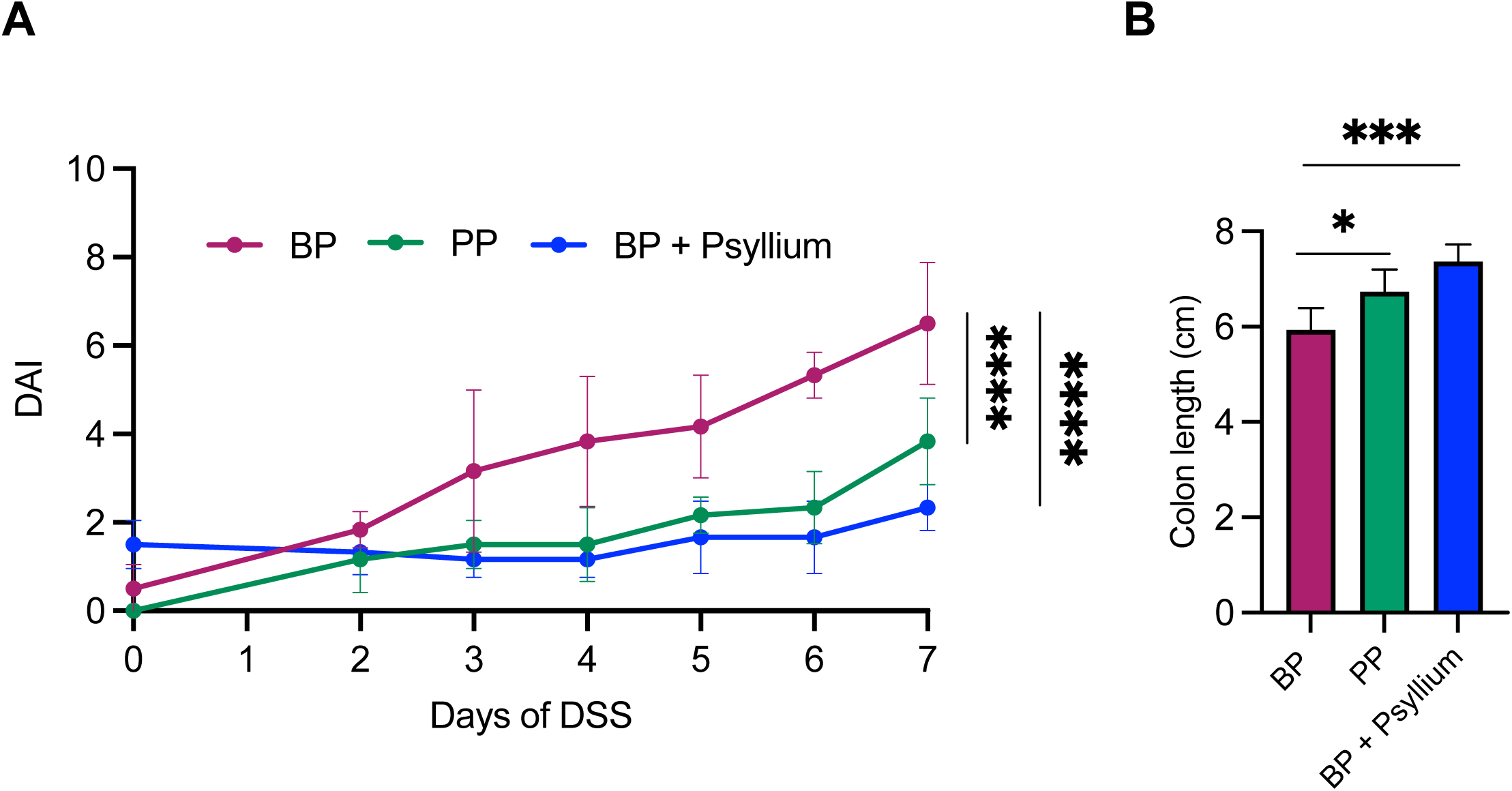
Psyllium supplementation protects against beef protein diet-induced worsening of DSS-induced colitis. WT specific pathogen free mice were fed beef protein (BP) or pea protein (BP) diets containing cellulose or BP diet containing psyllium (BP + Psyllium) for 7 days. Mice were then administered 1% dextran sodium sulfate (DSS) and Disease Activity Index (DAI) was measured over 7 days (A) and colon length was measured on day 7 (B). Bar plots show mean and standard deviation.

---

**Table S1.** Experimental Diet Formulations.

|  | Beef | Casein | Egg | Pea | Soy | Beef +<br>Psyllium |
| --- | --- | --- | --- | --- | --- | --- |
| Ingredient | gm | gm | gm | gm | gm | gm |
| Casein | 0 | 200 | 0 | 0 | 0 | 0 |
| Soy Protein, Supro 661 | 0 | 0 | 0 | 0 | 202.3 | 0 |
| Egg Whites | 0 | 0 | 210.9 | 0 | 0 | 0 |
| Beef Protein Isolate,<br>Sigma | 226.9 | 0 | 0 | 0 | 0 | 226.9 |
| Pea Protein, Pure Bulk | 0 | 0 | 0 | 209.1 | 0 | 0 |
| L-Cystine | 3 | 3 | 3 | 3 | 3 | 3 |
| Corn Starch | 398.486 | 397.486 | 389.286 | 398.486 | 395.486 | 385.329 |
| Maltodextrin 10 | 132 | 132 | 132 | 132 | 132 | 132 |
| Sucrose | 100 | 100 | 100 | 100 | 100 | 100 |
| Psyllium | 0 | 0 | 0 | 0 | 0 | 52.63 |
| Cellulose, BW200 | 50 | 50 | 50 | 50 | 50 | 0 |
| Soybean Oil | 70.8 | 70 | 70.2 | 51.8 | 63 | 70.8 |
| t-Butylhydroquinone | 0.014 | 0.014 | 0.014 | 0.014 | 0.014 | 0.014 |
| Mineral Mix S10022G | 0 | 0 | 0 | 0 | 0 | 0 |
| Mineral Mix S10001A | 17.5 | 17.5 | 17.5 | 17.5 | 17.5 | 17.5 |
| Calcium Phosphate,<br>Dibasic | 17.5 | 17.5 | 17.5 | 17.5 | 17.5 | 17.5 |
| Calcium Carbonate | 0 | 0 | 0 | 0 | 0 | 0 |
| Potassium Citrate,<br>Monohydrate | 0 | 0 | 0 | 0 | 0 | 0 |
| Potassium Phosphate,<br>Monobasic | 0 | 0 | 0 | 0 | 0 | 0 |
| Sodium Chloride | 0 | 20.2 | 12.9 | 19.3 | 19.2 | 0 |
| Vitamin Mix V10037 | 0 | 0 | 0 | 0 | 0 | 0 |
| Vitamin Mix V10001 | 10 | 10 | 10 | 10 | 10 | 10 |
| Biotin, 0.1% | 0 | 0 | 0.4 | 0 | 0 | 0 |
| Choline Bitartrate | 2.5 | 2.5 | 2.5 | 2.5 | 2.5 | 2.5 |
| Total | 1028.7 | 1020.2 | 1016.2 | 1011.2 | 1012.5 | 1018.2 |
| Protein (gm) | 177.0 | 177.0 | 177.0 | 177.0 | 177.0 | 177.0 |
| Carbohydrate (gm) | 640.5 | 640.5 | 640.5 | 640.5 | 640.5 | 640.5 |
| Fat (gm) | 71.0 | 71.0 | 71.0 | 71.0 | 71.0 | 71.0 |
| Fiber (gm) | 50.0 | 50.0 | 50.0 | 50.0 | 50.0 | 52.63 |
| Protein (kcal) | 708.1 | 708.0 | 708.0 | 707.9 | 707.9 | 708.1 |
| Carbohydrate (kcal) | 2561.9 | 2561.9 | 2561.9 | 2561.9 | 2561.8 | 2561.9 |
| Fat (kcal) | 639.2 | 638.6 | 639.4 | 639.3 | 639.3 | 639.2 |
| Total | 3909.3 | 3908.6 | 3909.3 | 3909.2 | 3909.0 | 3909.3 |
| Protein (gm%) | 17.2 | 17.3 | 17.4 | 17.5 | 17.5 | 17.2 |
| Carbohydrate (gm%) | 62.3 | 62.8 | 63.0 | 63.3 | 63.3 | 63.0 |
| Fat (gm%) | 6.9 | 7.0 | 7.0 | 7.0 | 7.0 | 6.9 |
| Protein (kcal%) | 18.1 | 18.1 | 18.1 | 18.1 | 18.1 | 18.1 |
| Carbohydrate (kcal%) | 65.5 | 65.5 | 65.5 | 65.5 | 65.5 | 65.5 |
| Fat (kcal%) | 16.4 | 16.3 | 16.4 | 16.4 | 16.4 | 16.4 |
| Sodium (g/kg) | 8.8 | 8.8 | 8.8 | 8.8 | 8.8 | 8.8 |
| kcal/gm | 3.8 | 3.8 | 3.8 | 3.9 | 3.9 | 3.8 |

**Table S2.**
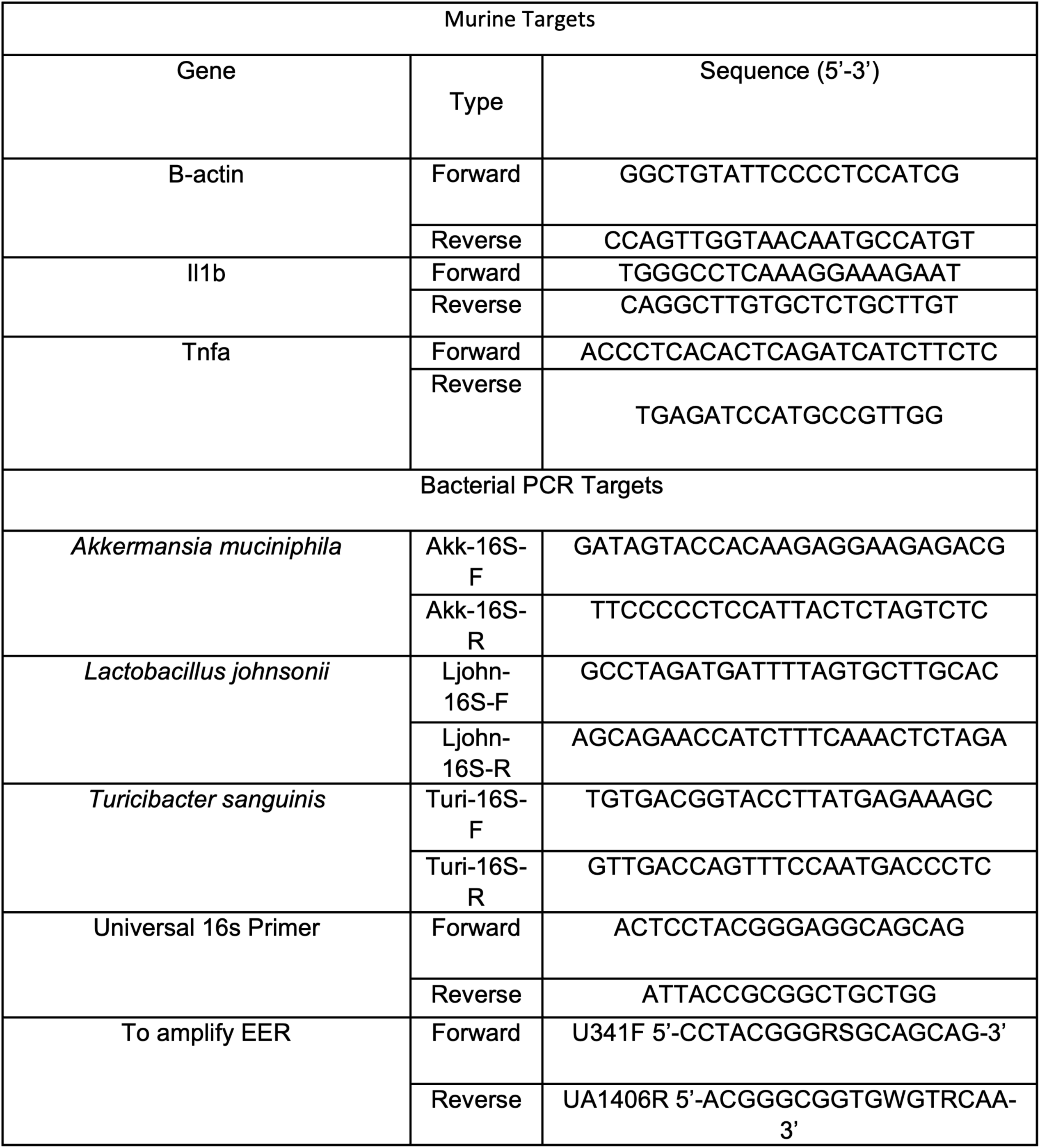
Primer Sequences.

## Notes

**Funding:** This work was supported by American Pulse Association and startup funds from the Stony Brook Cancer Center and Bahl Center for Metabolomics and Imaging (D.C.M.), NIH/NIDDK T32DK007737 (S.M.G., R.B.S.), P01DK094779 (R.B.S.), P40OD010995 (R.B.S.), P30DK034987 (R.B.S.), Crohn’s & Colitis Foundation Gnotobiotic Facility (R.B.S.), UNC Physician Scientist Training Program Fellowship Award (S.M.G.)

### Competing Interest Statement

The authors have declared no competing interest.

### Summary of Updates

Author name was misspelled in bioRxiv submission form as "R. Balfour Sator" and this has been corrected to "R. Balfour Sartor"

## References

1. Sartor RB, Wu GD. Roles for Intestinal Bacteria, Viruses, and Fungi in Pathogenesis of Inflammatory Bowel Diseases and Therapeutic Approaches. Gastroenterology. 2017;152(2):327–39.e4.

2. Perler BK, Friedman ES, Wu GD. The role of the gut microbiota in the relationship between diet and human health. Annual review of physiology. 2023;85:449–68.

3. Lee D, Albenberg L, Compher C, Baldassano R, Piccoli D, Lewis JD, et al. Diet in the Pathogenesis and Treatment of Inflammatory Bowel Diseases. Gastroenterology. 2015;148(6):1087–106.

4. Gleave A, Shah A, Tahir U, Blom J-J, Dong E, Patel A, et al. Using Diet to Treat Inflammatory Bowel Disease: A Systematic Review. Official journal of the American College of Gastroenterology | ACG. 2025;120(1).

5. Issokson K, Lee DY, Yarur AJ, Lewis JD, Suskind DL. The Role of Diet in Inflammatory Bowel Disease Onset, Disease Management, and Surgical Optimization. Official journal of the American College of Gastroenterology | ACG. 2025;120(1).

6. Hashash JG, Elkins J, Lewis JD, Binion DG. AGA Clinical Practice Update on Diet and Nutritional Therapies in Patients With Inflammatory Bowel Disease: Expert Review. Gastroenterology. 2024;166(3):521–32.

7. Kaplan GG, Ng SC. Understanding and Preventing the Global Increase of Inflammatory Bowel Disease. Gastroenterology. 2017;152(2):313–21.e2.

8. Kaplan GG, Windsor JW. The four epidemiological stages in the global evolution of inflammatory bowel disease. Nature Reviews Gastroenterology & Hepatology. 2021;18(1):56–66.

9. Kuenzig ME, Fung SG, Marderfeld L, Mak JWY, Kaplan GG, Ng SC, et al. Twenty-first Century Trends in the Global Epidemiology of Pediatric-Onset Inflammatory Bowel Disease: Systematic Review. Gastroenterology. 2022;162(4):1147–59.e4.

10. Daniel CR, Cross AJ, Koebnick C, Sinha R. Trends in meat consumption in the USA. Public Health Nutrition. 2011;14(4):575–83.

11. Al Hasan SM, Saulam J, Mikami F, Kanda K, Ngatu NR, Yokoi H, et al. Trends in per Capita Food and Protein Availability at the National Level of the Southeast Asian Countries: An Analysis of the FAO’s Food Balance Sheet Data from 1961 to 2018. Nutrients. 2022;14(3).

12. Desai MS, Seekatz AM, Koropatkin NM, Kamada N, Hickey CA, Wolter M, et al. A Dietary Fiber-Deprived Gut Microbiota Degrades the Colonic Mucus Barrier and Enhances Pathogen Susceptibility. Cell. 2016;167(5):1339–53.e21.

13. Hryckowian AJ, Van Treuren W, Smits SA, Davis NM, Gardner JO, Bouley DM, et al. Microbiota-accessible carbohydrates suppress Clostridium difficile infection in a murine model. Nature Microbiology. 2018;3(6):662–9.

14. Llewellyn SR, Britton GJ, Contijoch EJ, Vennaro OH, Mortha A, Colombel J-F, et al. Interactions Between Diet and the Intestinal Microbiota Alter Intestinal Permeability and Colitis Severity in Mice. Gastroenterology. 2018;154(4):1037–46.e2.

15. Kostovcikova K, Coufal S, Galanova N, Fajstova A, Hudcovic T, Kostovcik M, et al. Diet Rich in Animal Protein Promotes Pro-inflammatory Macrophage Response and Exacerbates Colitis in Mice. Front Immunol. 2019;10.

16. Riva A, Kuzyk O, Forsberg E, Siuzdak G, Pfann C, Herbold C, et al. A fiber-deprived diet disturbs the fine-scale spatial architecture of the murine colon microbiome. Nature Communications. 2019;10(1):4366.

17. Montrose DC, Nishiguchi R, Basu S, Staab HA, Zhou XK, Wang H, et al. Dietary Fructose Alters the Composition, Localization, and Metabolism of Gut Microbiota in Association With Worsening Colitis. Cellular and Molecular Gastroenterology and Hepatology. 2021;11(2):525–50.

18. Neumann M, Steimle A, Grant ET, Wolter M, Parrish A, Willieme S, et al. Deprivation of dietary fiber in specific-pathogen-free mice promotes susceptibility to the intestinal mucosal pathogen Citrobacter rodentium. Gut Microbes. 2021;13(1):1966263.

19. Wolter M, Steimle A, Parrish A, Zimmer J, Desai Mahesh S. Dietary Modulation Alters Susceptibility to Listeria monocytogenes and Salmonella Typhimurium with or without a Gut Microbiota. mSystems. 2021;6(6):e00717–21.

20. Shan Y, Lee M, Chang EB. The Gut Microbiome and Inflammatory Bowel Diseases. Annu Rev Med 2022;73:455–468.

21. Pereira GV, Boudaud M, Wolter M, Alexander C, De Sciscio A, Grant ET, et al. Opposing diet, microbiome, and metabolite mechanisms regulate inflammatory bowel disease in a genetically susceptible host. Cell Host & Microbe. 2024;32(4):527–42.e9.

22. Devkota S, Wang Y, Musch MW, Leone V, Fehlner-Peach H, Nadimpalli A, et al. Dietary-fat-induced taurocholic acid promotes pathobiont expansion and colitis in Il10-/- mice. Nature. 2012;487(7405):104–8.

23. Lloyd-Price J, Arze C, Ananthakrishnan AN, Schirmer M, Avila-Pacheco J, Poon TW, et al. Multi-omics of the gut microbial ecosystem in inflammatory bowel diseases. Nature. 2019;569(7758):655–62.

24. Sinha SR, Haileselassie Y, Nguyen LP, Tropini C, Wang M, Becker LS, et al. Dysbiosis-Induced Secondary Bile Acid Deficiency Promotes Intestinal Inflammation. Cell Host & Microbe. 2020;27(4):659–70.e5.

25. Yang ZH, Liu F, Zhu XR, Suo FY, Jia ZJ, Yao SK. Altered profiles of fecal bile acids correlate with gut microbiota and inflammatory responses in patients with ulcerative colitis. World J Gastroenterol. 2021:June 28;7(4):3609–29.

26. Joyce SA, Gahan CGM. Bile Acid Modifications at the Microbe-Host Interface: Potential for Nutraceutical and Pharmaceutical Interventions in Host Health. Annual review of food science and technology. 2016;7(1):313–33.

27. Gray SM, Moss AD, Herzog JW, Kashiwagi S, Liu B, Young JB, et al. Mouse adaptation of human inflammatory bowel diseases microbiota enhances colonization efficiency and alters microbiome aggressiveness depending on the recipient colonic inflammatory environment. Microbiome. 2024;12(1):147.

28. Png CW, Lindén SK, Gilshenan KS, Zoetendal EG, McSweeney CS, Sly LI, et al. Mucolytic Bacteria With Increased Prevalence in IBD Mucosa AugmentIn VitroUtilization of Mucin by Other Bacteria. Official journal of the American College of Gastroenterology | ACG. 2010;105(11).

29. Seregin SS, Golovchenko N, Schaf B, Chen J, Pudlo NA, Mitchell J, et al. NLRP6 Protects Il10-/- Mice from Colitis by Limiting Colonization of Akkermansia muciniphila. Cell Reports. 2017;19(4):733–45.

30. Sugihara K, Kitamoto S, Saraithong P, Nagao-Kitamoto H, Hoostal M, McCarthy C, et al. Mucolytic bacteria license pathobionts to acquire host-derived nutrients during dietary nutrient restriction. Cell Reports. 2022;40(3).

31. Hoek Kristen L, McClanahan Kathleen G, Latour Yvonne L, Shealy N, Piazuelo MB, Vallance Bruce A, et al. Turicibacterales protect mice from severe Citrobacter rodentium infection. Infect Immun. 2023;91(11):e00322–23.

32. Lynch JB, Gonzalez EL, Choy K, Faull KF, Jewell T, Arellano A, et al. Gut microbiota Turicibacter strains differentially modify bile acids and host lipids. Nature Communications. 2023;14(1):3669.

33. Zhang W, Wang J, Zhang D, Liu H, Wang S, Wang Y, et al. Complete Genome Sequencing and Comparative Genome Characterization of Lactobacillus johnsonii ZLJ010, a Potential Probiotic With Health-Promoting Properties. Frontiers in Genetics. 2019;10.

34. Jia D-J-C, Wang Q-W, Hu Y-Y, He J-M, Ge Q-W, Qi Y-D, et al. Lactobacillus johnsonii alleviates colitis by TLR1/2-STAT3 mediated CD206+ macrophagesIL-10 activation. Gut Microbes. 2022;14(1):2145843.

35. Kastl A, Zong W, Gershuni VM, Friedman ES, Tanes C, Boateng A, et al. Dietary fiber-based regulation of bile salt hydrolase activity in the gut microbiota and its relevance to human disease. Gut Microbes. 2022;14(1):2083417.

36. Jantchou P, Morois S, Clavel-Chapelon F, Boutron-Ruault M-C, Carbonnel F. Animal Protein Intake and Risk of Inflammatory Bowel Disease: The E3N Prospective Study. Official journal of the American College of Gastroenterology | ACG. 2010;105(10).

37. Jowett SL, Seal CJ, Pearce MS, Phillips E, Gregory W, Barton JR, et al. Influence of dietary factors on the clinical course of ulcerative colitis: a prospective cohort study. Gut. 2004;53(10):1479.

38. Lees CW, Gros B, Kyle JA, Plevris N, Derikx L, Constantine-Cooke N, et al. 477c HABITUAL MEAT INTAKE IS ASSOCIATED WITH INCREASED RISK OF DISEASE FLARE IN ULCERATIVE COLITIS: INITIAL RESULTS FROM THE PREDICCT STUDY. Gastroenterology. 2023;164(6):S-1571.

39. Jadhav A, Bajaj A, Xiao Y, Markandey M, Ahuja V, Kashyap PC. Role of Diet-Microbiome Interaction in Gastrointestinal Disorders and Strategies to Modulate Them with Microbiome-Targeted Therapies. Annu Rev Nutr. 2023;43:355–383.

40. Bibi S, De Sousa Moraes LF, Lebow N, Zhu M-J. Dietary Green Pea Protects against DSS-Induced Colitis in Mice Challenged with High-Fat Diet. Nutrients. 2017;9(5).

41. Ahn E, Jeong H, Kim E. Differential effects of various dietary proteins on dextran sulfate sodium-induced colitis in mice. Nutr Res Pract. 2022;16(6):700–15.

42. Chiang JYL, Ferrell JM. Bile Acids as Metabolic Regulators and Nutrient Sensors. Annu Rev Nutr. 2019;39:175–200.

43. Kubota H, Ishizawa M, Kodama M, Nagase Y, Kato S, Makishima M, et al. Vitamin D Receptor Mediates Attenuating Effect of Lithocholic Acid on Dextran Sulfate Sodium Induced Colitis in Mice. International Journal of Molecular Sciences. 2023;24(4).

44. Abdel-Razek EA-N, Mahmoud HM, Azouz AA. Management of ulcerative colitis by dichloroacetate: Impact on NFATC1/NLRP3/IL1B signaling based on bioinformatics analysis combined with in vivo experimental verification. Inflammopharmacology. 2024;32(1):667–82.

45. Wolter M, Grant ET, Boudaud M, Pudlo NA, Pereira GV, Eaton KA, et al. Diet-driven differential response of Akkermansia muciniphila modulates pathogen susceptibility. Molecular Systems Biology. 2024;20(6):596–625-.

46. Larabi AB, Masson HLP, Bäumler AJ. Bile acids as modulators of gut microbiota composition and function. Gut Microbes. 2023;15(1):2172671.

47. van Best N, Rolle-Kampczyk U, Schaap FG, Basic M, Olde Damink SWM, Bleich A, et al. Bile acids drive the newborn’s gut microbiota maturation. Nature Communications. 2020;11(1):3692.

48. Islam KBMS, Fukiya S, Hagio M, Fujii N, Ishizuka S, Ooka T, et al. Bile Acid Is a Host Factor That Regulates the Composition of the Cecal Microbiota in Rats. Gastroenterology. 2011;141(5):1773–81.

49. Tian YA-O, Gui W, Koo IA-O, Smith PB, Allman EA-O, Nichols RG, et al. The microbiome modulating activity of bile acids. Gut Microbes. 2020(Mar 5;11(4):979–996.).

50. Bretin A, Zou J, San Yeoh B, Ngo VL, Winer S, Winer DA, et al. Psyllium Fiber Protects Against Colitis Via Activation of Bile Acid Sensor Farnesoid X Receptor. Cellular and Molecular Gastroenterology and Hepatology. 2023;15(6):1421–42.

51. Sartor RB. Mechanisms of Disease: pathogenesis of Crohn’s disease and ulcerative colitis. Nature Clinical Practice Gastroenterology & Hepatology. 2006;3(7):390–407.

52. Yang C, Merlin D. Unveiling Colitis: A Journey through the Dextran Sodium Sulfate-induced Model. Inflammatory Bowel Diseases. 2024;30(5):844–53.

53. Faith JJ, Ahern PP, Ridaura VK, Cheng J, Gordon JI. Identifying Gut Microbe–Host Phenotype Relationships Using Combinatorial Communities in Gnotobiotic Mice. Science Translational Medicine. 2014;6(220):220ra11–ra11.

54. Darfeuille-Michaud A, Neut C, Barnich N, Lederman E, Di Martino P, Desreumaux P, et al. Presence of adherent Escherichia coli strains in ileal mucosa of patients with Crohn’s disease. Gastroenterology. 1998;115(6):1405–13.

55. Eun CS, Mishima Y, Wohlgemuth S, Liu B, Bower M, Carroll IM, et al. Induction of bacterial antigen-specific colitis by a simplified human microbiota consortium in gnotobiotic interleukin-10-/- mice. Infect Immun. 2014;82(6):2239–46.

56. Remke M, Groll T, Metzler T, Urbauer E, Kövilein J, Schnalzger T, et al. Histomorphological scoring of murine colitis models: A practical guide for the evaluation of colitis and colitis-associated cancer. Experimental and Molecular Pathology. 2024;140:104938.

57. Rath HC, Herfarth HH, Ikeda JS, Grenther WB, Hamm TE, Jr., Balish E, et al. Normal luminal bacteria, especially Bacteroides species, mediate chronic colitis, gastritis, and arthritis in HLA-B27/human beta2 microglobulin transgenic rats. J Clin Invest. 1996;98(4):945–53.

58. Chassaing B, Srinivasan G, Delgado MA, Young AN, Gewirtz AT, Vijay-Kumar M. Fecal Lipocalin 2, a Sensitive and Broadly Dynamic Non-Invasive Biomarker for Intestinal Inflammation. PLoS One. 2012;7(9):e44328.

59. Bai JDK, Saha S, Wood MC, Chen B, Li J, Dow LE, et al. Serine Supports Epithelial and Immune Cell Function in Colitis. The American Journal of Pathology. 2024;194(6):927–40.

60. Derrien M, van Baarlen P, Hooiveld G, Norin E, Muller M, de Vos W. Modulation of Mucosal Immune Response, Tolerance, and Proliferation in Mice Colonized by the Mucin-Degrader Akkermansia muciniphila. Front Microbiol. 2011;2.

61. Chetnik K, Benedetti E, Gomari DP, Schweickart A, Batra R, Buyukozkan M, et al. maplet: an extensible R toolbox for modular and reproducible metabolomics pipelines. Bioinformatics. 2022;38(4):1168–70.

